# Proteomic cellular signatures of kinase inhibitor-induced cardiotoxicity: Mount Sinai DToxS LINCS Center Dataset

**DOI:** 10.1101/2020.02.26.966606

**Authors:** Yuguang Xiong, Tong Liu, Tong Chen, Jens Hansen, Bin Hu, Yibang Chen, Gomathi Jayaraman, Stephan Schürer, Dusica Vidovic, Joseph Goldfarb, Eric A. Sobie, Marc R. Birtwistle, Ravi Iyengar, Hong Li, Evren U. Azeloglu

## Abstract

The Drug Toxicity Signature Generation Center (DToxS) at the Icahn School of Medicine at Mount Sinai is one of the centers of the NIH Library of Integrated Network-Based Cellular Signatures (LINCS) program. A key aim of DToxS is to generate both proteomic and transcriptomic signatures that can predict adverse effects, especially cardiotoxicity, of kinase inhibitors approved by the Food and Drug Administration. Towards this goal, high throughput shot-gun proteomics experiments (317 cell line/drug combinations + 64 control lysates) have been conducted at the Center for Advanced Proteomics Research at Rutgers University - New Jersey Medical School. Using computational network analyses, these proteomic data can be integrated with transcriptomic signatures generated in tandem to identify cellular signatures of cardiotoxicity that may predict kinase inhibitor-induced toxicity and possible mitigation. Both raw and processed proteomics data have been carefully screened for quality and made publicly available via the PRIDE database. As such, this broad protein kinase inhibitor-stimulated cardiomyocyte proteomic data and signature set is valuable for the prediction of drug toxicities.

**Links to: Metadata Tables:** 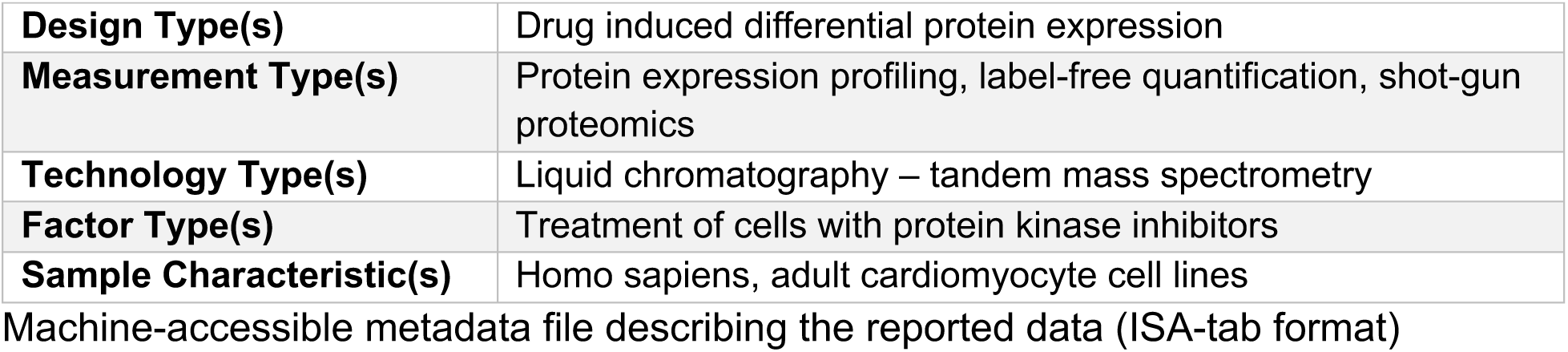

## Background & Summary

Protein kinase inhibitors (KIs) are a class of targeted therapeutics that are being increasingly used in the treatments of cancer ^1^. Their use and development have been accelerated in the recent years as they could target tumors more effectively than most other chemotherapeutics and their mechanisms of actions are well defined. In many cases, however, their intended or off-target kinases serve key biological roles, which when blocked, lead to severe adverse effects ^2^. One of the major adverse effects that lead to KI discontinuation of treatment is cardiotoxicity ^3^. As a part of the NIH Library of Integrated Cellular Signatures (LINCS) program ^4^, the main goal of the Mount Sinai Drug Toxicity Signature Generation Center (DToxS) is to better understand mechanisms of KI-associated cardiotoxicity by constructing cellular signatures of drug effects. Since different omics assays show varied sensitivities ^5^ and offer complementary molecular information on the cellular phenotypic state ^6^, in order to develop a comprehensive understanding of the cellular responses to KIs, we use network analyses ^7^ to combine differential expression of genes and gene products in human cardiomyocytes treated by FDA approved KIs (**Table 1**), analyzed using both transcriptomic and proteomic methods. In this dataset, we present the proteomic portion of the effects of KIs on human cardiomyocytes.

**Table 1.** Drug metadata. Purity was verified for each lot number. The concentration used for each drug is equivalent to the clinically-observed median peak plasma concentration as per the cited reference.

| Drug | Acronym | Drug Form | Purity | Concentration ($\mu\text{M}$) | Reference |
| --- | --- | --- | --- | --- | --- |
| **Afatinib** | AFA |  | 98.8 | 0.05 | 27,28 |
| **Axitinib** | AXI |  | 98.0 | 0.2 | 29,30 |
| **Bosutinib** | BOS |  | 98.0 | 0.1 | 31 |
| **Cabozantinib** | CAB |  | 99.1 | 2 | 32 |
| **Dabrafenib** | DAB |  | 99.61 | 2.5 | 33,34 |
| **Dasatinib** | DAS |  | 98.36 | 0.1 | 35 |
| **Erlotinib** | ERL |  | 99.32 | 3 | 36,37 |
| **Gefitinib** | GEF |  | 99.25 | 1 | 38 |
| **Imatinib** | IMA |  | 99.48 | 5 | 39 |
| **Lapatinib** | LAP |  | 99.0 | 2 | 40 |
| **Nilotinib** | NIL |  | 99.77 | 3 | 41,42 |
| **Pazopanib** | PAZ |  | 96.13 | 10 | 43 |
| **Ponatinib** | PON |  | 97.29 | 0.1 | 44 |
| **Regorafenib** | REG |  | 99.58 | 1 | 45 |
| **Ruxolitinib** | RUX |  | 99.38 | 1 | 46,47 |
| **Sorafenib** | SOR | Tosylate | 99.73 | 0.5 | 46,47 |
| **Sunitinib** | SUN | Malate | 97.13 | 1 | 48,49 |
| **Trametinib** | TRA |  | 99.24 | 0.1 | 50 |
| **Tofacitinib** | TOF | Citrate | 99.85 | 1 | 50 |
| **Vandetanib** | VAN |  | 99.73 | 0.333 | 51 |
| **Vemurafenib** | VEM |  | 98.08 | 2 | 52 |

Quantitative proteomics technologies have evolved to become increasingly effective at identifying differentially expressed proteins among diverse experimental conditions. Currently, two broad strategies are widely used for large-scale quantitative proteomics studies: stable isotope label-based ^8,9^ and label-free ^10,11^ quantification (LFQ) strategies. Each strategy entails trade-offs of strengths and limitations. Stable isotope label-based strategies involve labeling proteins or peptides with amino acids or chemical tags that contain stable isotopes (*e.g.* ^2^H, ^13^C, ^15^N or ^18^O, *etc.)*, which provide mass shift features that are differentiated in high resolution mass spectrometers for the quantification of proteins pooled from up to a dozen biological conditions. Examples include Isobaric Tag for Relative and Absolute Quantitation (iTRAQ), Tandem Mass Tag (TMT) and Stable Isotope Labeling with Amino acids in Cell culture (SILAC). Key benefits of the label-based proteomics strategy over LFQ methods include the minimization of experimental variations and the maximization of protein quantitation precisions ^12,13^. In contrast, two key factors limit the wide applications of the label-based strategies for clinically relevant proteomics studies of hundreds of samples: commercially available stable isotope proteomics reagents can currently accommodate the simultaneous quantification of up to 11 samples; and the costs for label-based proteomics reagents are prohibitively expensive for analyzing hundreds of samples. Compared to label-based proteomics strategies, LFQ methods do not involve expensive labeling reagents, thus they can be used for clinically relevant studies of hundreds and thousands of samples, if proper controls and quality guidelines are followed. The most widely used LFQ strategies utilize either the peptide mass spectral ion intensities or tandem mass spectral counts to approximate the abundances of proteins and compare them across many samples. Also, LFQ methods offer additional benefits over label-based strategies, including quantification with wider dynamic ranges and more flexibility in study design ^14-16^. When compared to label-based strategies, the trade-offs for the LFQ methods are well known: results may contain larger variability in quantification and smaller number of proteins quantified. Larger quantification variability can originate from the fact that in a LFQ workflow, each sample is analyzed independently, including proteolytic digestion and LC/MS/MS analysis, which can introduce additive technical variations at each experimental step. These analytical inconsistencies can be minimized by omitting multi-dimensional peptide fractionations; however, while LFQ sample throughput can be increased through this approach, it should be noted that this approach can also lead to a smaller number of identified proteins.

In this DToxS dataset, we’ve chosen the LFQ method in order to economically compare the proteomics signatures from over 300 samples that have been collected over an extended period of time, some with limited protein yields. The transcriptome of these samples have been previously assessed using the 3’ digital gene expression RNA sequencing (RNAseq) ^17^; to ensure both proteomics and RNAseq data can be compared from the same drug-treated cells (see example use case below), we’ve optimized a method to extract proteins from the samples after RNA extraction (**Fig. 1**). All the sample preparation, analysis procedures and data have gone through careful quality control steps to ensure reproducibility (**Fig. 2**); standard operating procedures and data descriptors are publicly available in the DToxS website (www.dtoxs.org) and both raw and MaxQuant-analyzed data are publicly available via PRIDE (with the proteome exchange ID of PXD014791) as well as the LINCS Project data portals (http://lincsportal.ccs.miami.edu/dcic-portal/). We think that this large KI-induced proteomics dataset will be a valuable resource for the cancer, cardiology and drug development communities.

**Figure 1.**
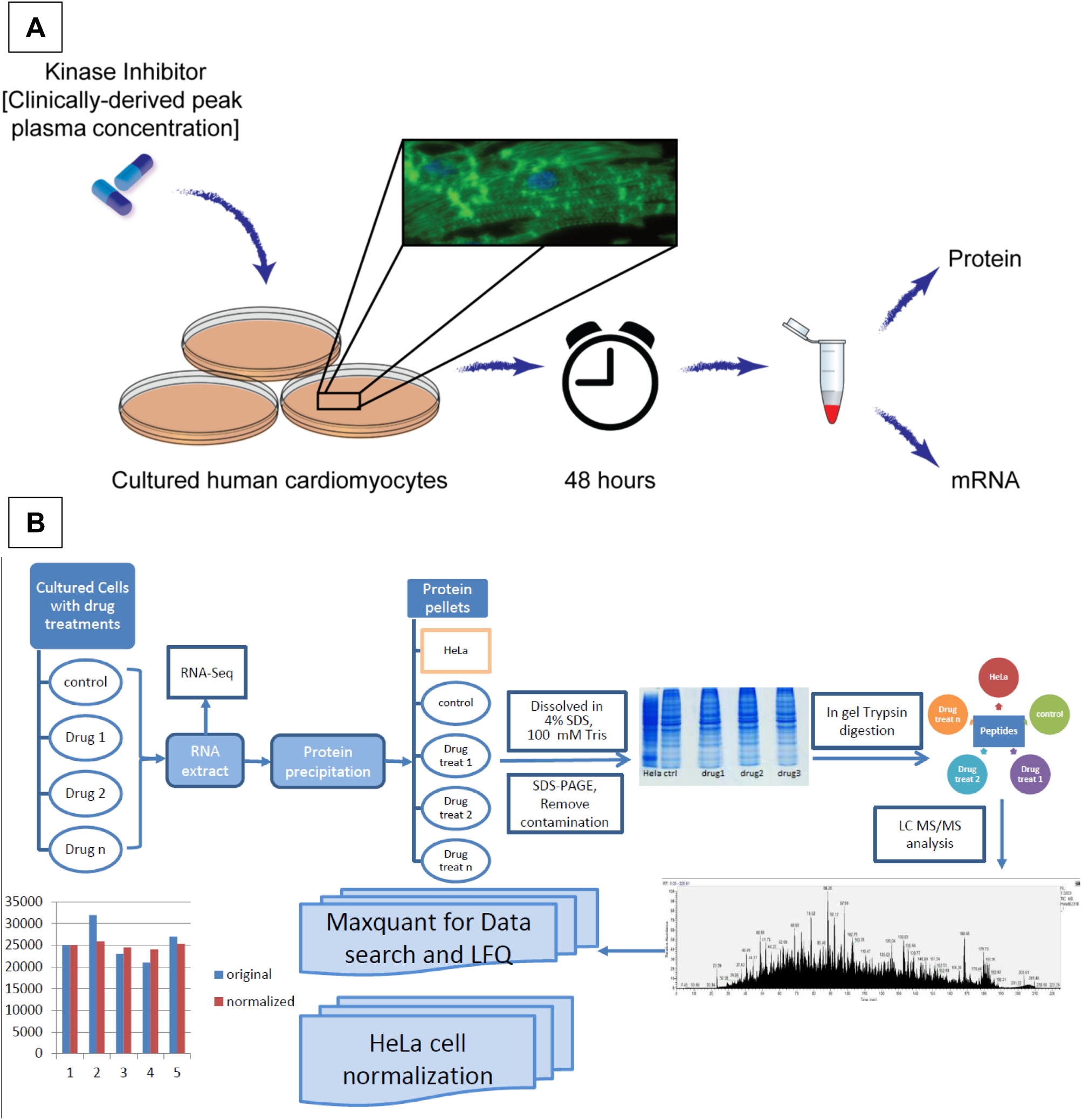
LFQ proteomics workflow for the DToxS Study. **(A)** Drug and cell treatment design and experimental workflow. **(B)** After drug treatment, RNA were isolated from the cells, and the remaining proteins were recovered via protein precipitation. The protein amounts were carefully estimated from their SDS-PAGE staining intensities as measured relative to a HeLa cell lysate standard. Two μg of protein from each sample were analyzed by LC-MS/MS, and the resulting proteins were quantified by a LFQ approach using MaxQuant.

**Figure 2.**
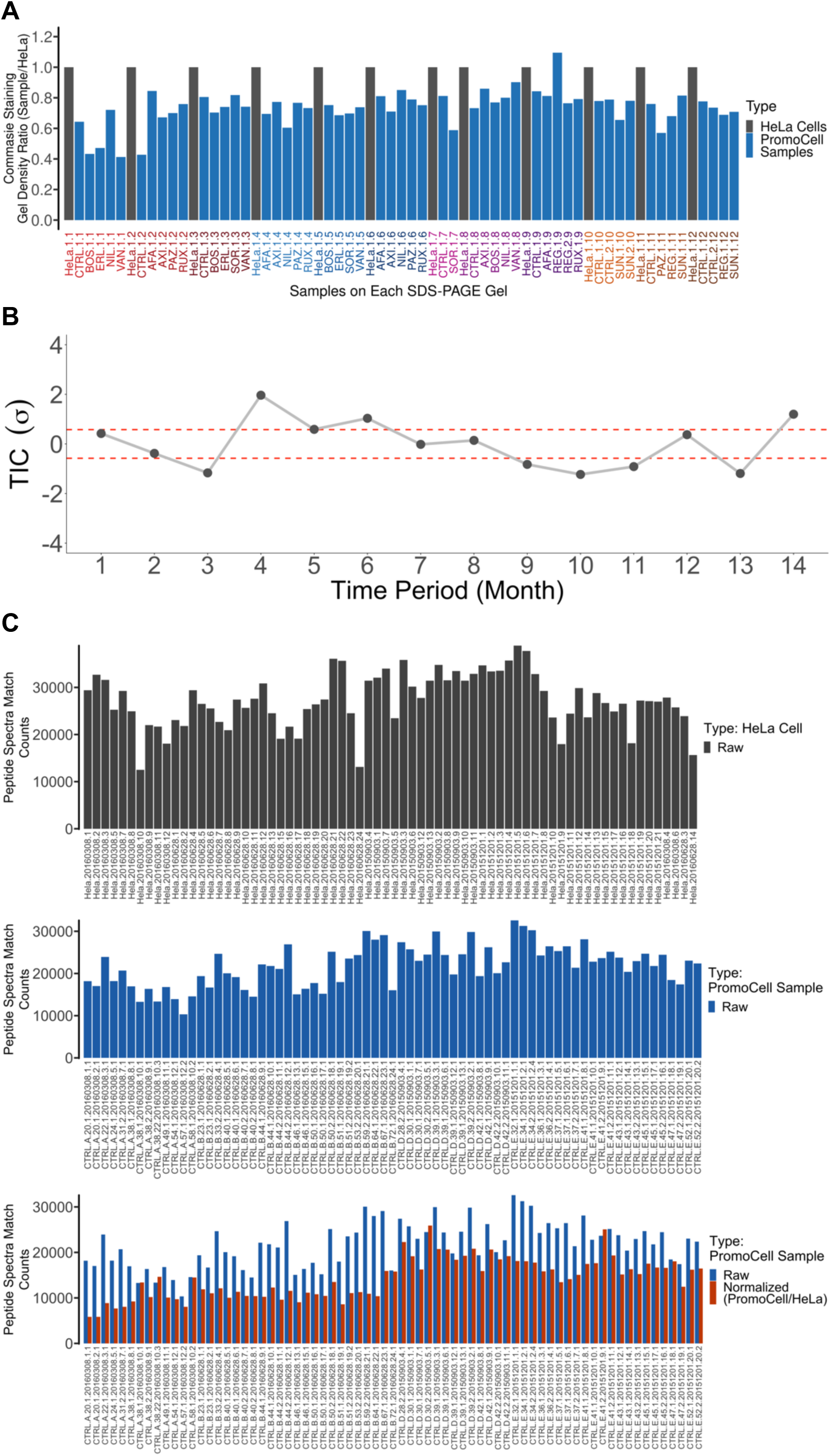
MS/MS quality control for the DToxS Study. A series of quality control steps have been implemented to obtain high quality data for this study. **(A)** A gel-based method developed for protein estimation. In each SDS-PAGE gel, 25 μg of HeLa cell protein extract was run as a reference to estimate the total protein amount of each sample (blue verticle lines). Based on the density of the CBB stain of each lane in relation to the HeLa stain density, the protein amount can be calculated (see examples in Supplementary Table 1). After the tryptic digestions, the resulting peptides were diluted into 0.5 μg/ml for LC-MS/MS analysis (orange verticle lines), based on the estimated protein amount in each sample. **(B)** A commercial HeLa digest used for LC-MS/MS quality control. Two hundred ng of HeLa protein digest was analyzed regulary to ensure good LC-MS/MS instrument performance. During the time period of the DToxS LINCS data acquisition, the total ion current (TIC) of a representative HeLa cell digestion from each month is plotted monthly, indicating consistent LC-MS signal response from the commercial HeLa digest. **(C)** Deep proteome coverage indicates good instrument sensitivity. The total protein groups identified from each cell line in this study are shown, with ∼7,000 proteins identified per cell line. **(D)** The HeLa protein digest from (A) were used LFQ normalization. Based on the MS1 LFQ counts of the HeLa cells from each gel (top panel), a normalization factor was calculated and applied to the MS1 counts of the samples (the control cell data before normalization is shown in the middle panel). After the HeLa cell normalization, the LFQ variation was dramatically reduced (lower panel).

## Methods

### Cell culture, drug treatments and RNA extraction

Detailed materials and methods for all the experiments in this study can be accessed as version- and quality-controlled standard operating procedures on DToxS.org. Briefly, four commercially available cell lines of primary adult human cardiomyocytes were purchased from PromoCell GmbH (Cat #: C12810; Heidelberg, Germany), expanded and differentiated under serum-free conditions for 28 days per manufacturer’s instructions. The four cell lines used in this study (Lot #: 3042901.2, 4031101.3, 2082801.2, 2120301.2) were isolated from two Caucasian male and two Caucasian female subjects, aged 54, 62, 61, 56, respectively (See cell lines A, B, D, E in **Table 2**). Although these cell lines were originally derived from the heart and have many cardiomyocyte properties, they are non-excitable or contractile; hence, we refer to them as cardiomyocyte-like throughout the dataset. Briefly, cells were subcultured at 37 °C under 5% CO_2_ with the manufacturer-supplied and serum-supplemented growth media (Cat #: C22270, C39275) up until passage four. Once they reached 100% confluence, cells were trypsinized, counted, and replated in serum-free growth media (Cat #: C22270), at a concentration of 40,000 cells/cm^2^ in 60 mm tissue culture dishes. Cells were then differentiated under serum-free conditions for 28 days, whereby 50% of the media was replenished every other day. All experiments were performed on sixth or earlier passage cells. After 28 days of differentiation, cells were treated with individual KIs for 48 hours at a concentration equivalent to clinically-observed median peak plasma concentration (see **Table 1** for additional drug metadata). After 48 hours of KI treatment (see the drug treatment schedule in **Table 2**), cells were lysed on ice using TRIzol (Thermo Fisher, Cat #: 15596026) for five minutes, scraped off the dish, and mixed. TRIzol lysates were mixed with chloroform according to manufacturer’s instructions and the RNA-protein fractions were separated. From these organic partitions, RNA samples were processed, sequenced and analyzed as previously reported ^17^.

**Table 2.** Cellular metadata.

| Cell Name | Vendor | Cat No | Lot No | Race | Sex | Age | Disease |
| --- | --- | --- | --- | --- | --- | --- | --- |
| PMC-A | PromoCell | C-12810 | 3042901.2 | Caucasian | Male | 54 | None |
| PMC-B | PromoCell | C-12810 | 4031101.3 | Caucasian | Female | 62 | None |
| PMC-D | PromoCell | C-12810 | 2082801.2 | Caucasian | Female | 61 | None |
| PMC-E | PromoCell | C-12810 | 2120301.2 | Caucasian | Male | 56 | None |

**Table 3.** Summary of MaxQuant analytics.

| **Group** | **PMC-A** | **PMC-B** | **PMC-D** | **PMC-E** |
| --- | --- | --- | --- | --- |
| **Total Identified Protein Groups** | 6,803 | 6,968 | 6,945 | 7,021 |
| **Total Identified Peptides** | 1,599,186 | 3,266,823 | 2,023,891 | 3,227,847 |
| **Total Unique Peptides** | 1,090,438 | 2,224,679 | 1,423,505 | 2,252,728 |
| **Median LFQ Intensity** | 14,430,000 | 15,755,000 | 13,306,000 | 12,503,000 |

### Protein extraction

After isolating RNA, the remaining TRIzol solutions were processed for protein extractions. Each TRIzol extract were first separated into two 1.5 mL Eppendorf tubes; each tube contained ∼600 μl of the TRIzol chloroform and methanol solvent mixture. Then to each tube, 900 μl of pre-chilled cold acetone was added and mixed well. Five hundred μL of the mixture was transferred from each tube to a third new tube, and 500 μL pre-chilled −20°C acetone was added to each of the three tubes and mixed well. All three tubes were put on ice for at least 12 hours inside a cold room, and centrifuged at 16,000 rpm (∼23,469 g) for 30 min, in a Beckman Coulter Allegra 64R centrifuge. After centrifugation, the supernatants were removed and 1 mL of pre-chilled acetone were added to each tube to wash the protein pellets. The protein pellets were sonicated for five bursts at level-3, repeated once. The protein solutions were placed on ice for 4 hours and pelleted *via* centrifugation as described above. The wash procedure for each protein pellet was repeated twice for a total of three times. The final protein pellet was washed with 50 μL of pre-chilled acetone and centrifuged at 16,000 rpm for 10 min, and the remaining pellet was air dried at room temperature for 10 to 15 minutes. The protein pellet was then resolubilized with a vortex mixer, with 70 μl of 8 M urea (Fisher Cat #: U15500) in 50 mM ammonium bicarbonate (Sigma Cat#: 09830-500G), and stored at −80°C before proteomics analysis. See additional protocol details at https://martip03.u.hpc.mssm.edu/sop.php.

### Sample preparation for proteomics analysis (Fig. 2)

The protein solutions may contain TRIzol and other low mass contaminants from the RNA extraction buffer, which can confound accurate protein estimation *via* conventional biochemical assays. Consequently, we ran the SDS-PAGE gels to remove the low mass contaminants from the high mass proteins, and estimated the protein amounts *via* the stain intensities from the gels stained with the Coomassie brilliant blue (CBB) dyes (see examples in **Supplementary Fig. 1**). In each gel, a HeLa cell extract (Pierce, Cat # 88329, 25 μg) was run as the normalization reference, along with the proteins isolated from the drug-treated cell lines. After Coomassie blue staining of the gels, the density of each sample lane was measured using the ImageJ software (NIH), and the protein amount for each drug-treated sample was calculated using the following equation: protein (μg) = HeLa protein 25 μg × (Sample density / HeLa density). The protein quantity information was later used to dilute the tryptic peptides into 0.5 µg protein equivalent per µl for the LC-MS/MS analysis (**Supplementary Table 1**).

The in-gel digestions of proteins were performed following a procedure similar to the one described by the Mann lab ^18^. In brief, the entire gel lane of each sample was excised into ∼ 1 cm^2^ gel blocks, and washed four times with 10 mL each of a Wash Buffer containing 30% acetonitrile (ACN) and 70% of 100 mM NH_4_HCO_3_ to remove the contaminants from the RNA extraction buffer. Subsequently, one mL of 25 mM dithiothreitol (DTT) solution was added to the gel blocks for disulfide reduction at 55°C for 30 min, and 1 mL of 50 mM iodoacetamide solution was then added for thiol alkylation at 37°C in the dark for 30 min. After alkylation, the gel blocks were dehydrated with 9 mL of ACN to remove both DTT and iodoacetamide. For in-gel trypsin digestion, one mL of trypsin solution (5 µg/mL in 50 mM NH_4_HCO_3_) was added into each sample, and incubated at 37°C for 16 h. Resulting peptides were extracted, desalted with Pierce C_18_ spin columns (Thermo Scientific) based on the manufacturer’s protocol and concentrated in a Speed Vac prior to LC-MS/MS analysis.

### LC-MS/MS analysis

According to the gel-based protein amount estimates described above, the peptides from each sample were resuspended in Solvent A (2% ACN in 0.1% formic acid (FA)) at a final concentration of 0.5 µg/µL (see examples in **Supplementary Table 1**). Two microgram of peptides from each sample was subjected to LC-MS/MS analysis on a Q Exactive Mass Spectrometer coupled with an UltiMate 3000 RSLCnano Duo LC system (Thermo Scientific). The peptides were first loaded onto a trapping column (Acclaim PepMap 100 C_18_ trap column 75 µm × 2 cm, 3 µm, 100 Å) and then separated on an Acclaim PepMap C_18_ column (75 µm × 50 cm, 2 µm, 100 Å), using a 4-h binary gradient from 2-100% of Solvent B (85% ACN in 0.1% FA), at a flow rate of 250 nL/min. The eluted peptides were directly introduced into the MS system for data-dependent MS/MS analysis in the positive ion mode. The MS full scans were acquired in an m/z range of 400 to 1750, with the AGC value specified at 3E6, the injection time set at 100 ms, and in the profile mode. The resolution of the full MS scan was set to 140,000 at m/z 400. Following each full MS scan, 15 most intense ions with charge states between 2^+^ to 5^+^ were selected within an isolation window of 2 m/z for the subsequent MS/MS analysis. The AGC of MS/MS analysis was set to 5E4 and the dynamic exclusion was 45 s. The peptide ions were fragmented using higher energy collision dissociation at a NCE of 27. A total 317 LC-MS/MS raw files were obtained for the proteins isolated across four different cell lines, with a total 62 different types of drug treatment conditions. Prior to running each set of drug-treated samples, a HeLa cell digest (Pierce, Cat # 88329) was run as a reference to quality control the LC-MS/MS system.

### MaxQuant for protein identification and quantification

In order to evaluate the quality of the data and compare the quantitative proteomic signatures among the drug-treated samples across different cell lines, the entire 381 raw LC-MS/MS dataset (317 samples and 64 HeLa cell controls, ∼ 825 GB of data) were submitted for database search using the Andromeda search engine on the MaxQuant platform (Version 1.6.0.13). The raw data files were loaded with “No fractions” option selected. Trypsin was selected as enzyme with two miss cleavages. Methionine oxidation (+15.9949 Da) and protein N-terminal acetylation (+42.0106 Da) were selected as various modifications and cysteine carbamidomethyl modification (+57.0215 Da) was set as a fixed modification. Initial search peptide mass tolerance was set to 20 ppm, and the main search peptide mass tolerance was set to 4.5 ppm. LFQ was selected for label-free quantification, with the minimal LFQ ratio count set at 2. The MS/MS spectra were searched against both UniProt human FASTA database (downloaded from https://www.uniprot.org/proteomes/UP000005640 with the last modification date of 10/22/2018, containing 73,101 human protein sequences) and the MaxQuant default contaminants FASTA database (containing 245 protein sequences). Match between runs was selected for maximize the protein identification and quantitation with a match time window of 0.7 min and an alignment time window of 20 min. The protein false discovery rate (FDR) was estimated using the decoy databases containing reversed sequences of the original proteins. Proteins identified with both protein and peptide FDRs at or less than 1% were included in the final results for the subsequent analyses.

## Data Records

All the raw data described in this study have been uploaded to ProteomeXchange website (http://www.proteomexchange.org/) with the project accession PXD014791, and they are freely available for the research community. In addition, the processed higher level data is available through LINCS Data Portal (http://lincsportal.ccs.miami.edu/dcic-portal/). Data include metadata (Data Citation 1), raw files of LC-MS/MS analysis (Data Citation 2), protein FASTA database (Data Citation 3), and protein identification and quantitation results from the MaxQuant (Data Citation 4).

## Technical Validation

We’ve developed a 3-tiered quality control strategy to ensure the high throughput production of deep and reproducible DToxS proteomics datasets. This strategy has enabled us to control the qualities at the protein sample level, the LC-MS/MS instrument performance level and the overall sample processing variations.

### Quality validation of the protein samples for LC-MS/MS

Due to both the cost constrains of treating the human-derived cardiomyocyte-like cells with KI drugs and the scientific necessity to compare proteomics data with RNAseq data from the same samples, the same drug-treated cells were used to analyze protein expression, after the RNA extraction. The challenges from this approach included occasionally uneven and poor protein yields, and imprecise protein concentration estimates with the established Bradford or BCA assays, due to the confounding components in the RNA extraction buffers. In order to rigorously compare drug effects with the given high sample-to-sample variability in experimental processing, we performed an additional SDS-PAGE in-gel quantification step with a consistent positive control sample. Seventy SDS-PAGE gels were run to increase the estimation accuracy of the proteins derived from the cells (see examples in **Supplementary Fig. 1**); in each gel, 25 μg of HeLa cell protein lysate was included in the first lane, as the reference for both accurate protein estimation and efficient in-gel digestion, and later for LC-MS/MS data normalization. In brief, the densities of the CBB for each KI-treated cells in each gel lane were recorded (see examples in **Supplementary Table 2**). The protein amounts from each sample were extrapolated by comparing its CBB density to that of the 25 μg of the HeLa proteins; the estimated protein amounts were later used to obtain 0.5 µg/µL equivalent of the tryptic peptides from each sample for LC-MS/MS analysis (see examples in **Fig. 2A**). For tryptic digestion, the drug-treated samples were digested along with the HeLa cell proteins separated on the same SDS-PAGE gels, which were later used for LC-MS/MS data normalization as well.

### Quality validation of the LC-MS/MS system

To ensure that LC-MS/MS system could deliver overall satisfactory performance, 200 ng of the commercial HeLa cell digest was run prior to each gel batch of the biological samples. To confirm well-behaved LC separations, MS1 ions for select peptides ions were carefully monitored for retention time shrifts and peak widths. In order to minimize the run-to-run carry-over of peptides, a duo-LC system was configurated in parallel for this study, so while one LC column was used for peptides separation and LC-MS/MS analysis, the other LC column could be washed to remove any residual peptides. For the over LC-MS/MS quality control, the system is capable of identifying > 3,800 unique proteins and > 14,000 unique peptides from the 200 ng of the commercial HeLa cell digest. Indeed, monthly total ion current (TIC) analysis of the HeLa digest indicates consistent LC-MS/MS system performance for LFQ quantification (**Fig. 2B**). Qualitatively, the LC-MS/MS system enabled the deep proteome coverages (∼ 7,000 unique protein groups/cell line) of all four DToxS cell lines (**Fig. 2C** and **Table 2**). Total 381 raw LC-MS/MS files, including the files for 317 drug treat-cells and 64 HeLa cells, were submitted to the MaxQuant analysis (**Table 2**). In total, 7,193 protein groups were identified with the FDR of less than 1%, among the 4 different cell lines, which were mapped to 5,734 genes. Given no multi-dimensional LC fractionation was performed, these deep proteome coverages confirm the effectiveness of the LC-MS/MS quality control procedures used for this study.

### Quality control of the quantitative data analysis

To minimize the LC-MS/MS quantification noise derived from the sample preparation steps, the HeLa proteins used for the in-gel protein estimation were digested along with the drug-treated samples from the same gels. Thus, these HeLa cell protein digests were used to control for the trypsin digestion efficiencies, LC/MS/MS variations and for the quantitative normalization of the LFQ data. For example, the TIC analyses indicates sample-to-samples variations among both HeLa and control cell (no drug treatment) digests (**Fig. 2D**, upper and middle panel, respectively); with HeLa TIC signal normalization, the variations in the control cell digests were dramatically reduced (**Fig. 2D**, lower panel).

Overall, our method for protein count normalization showed remarkable reproducibility (**Fig 3A**), which can be assessed using Pearson correlation across two technical replicates that were analyzed on different days from two different HeLa signal normalization protocols (i.e., different SDS-PAGE gels). This was consistent for all technical replicates in this dataset (**Supplementary Fig 2**). We note that while depth of protein quantification was high for each replicate, overlap of detected proteins between different biological replicates was consistently high for all cardiomyocyte-like cell lines (**Fig 3B**), suggesting that our experimental pipeline was robust in identification of proteomic signatures. The quantification strategy was also successful in reproducible identification of subtle proteomic differences between individual cardiomyocyte cell lines PMC-A, B, D and E, which clustered consistently across most biological replicates (**Fig 3B**). Furthermore, we used overlap analysis, to show that most of the identified proteins were common to either all four cell lines (3,784 or 66%) or detected in at least three out of four lines (4,499 or 78.5%) (**Fig 4**).

**Figure 3.**
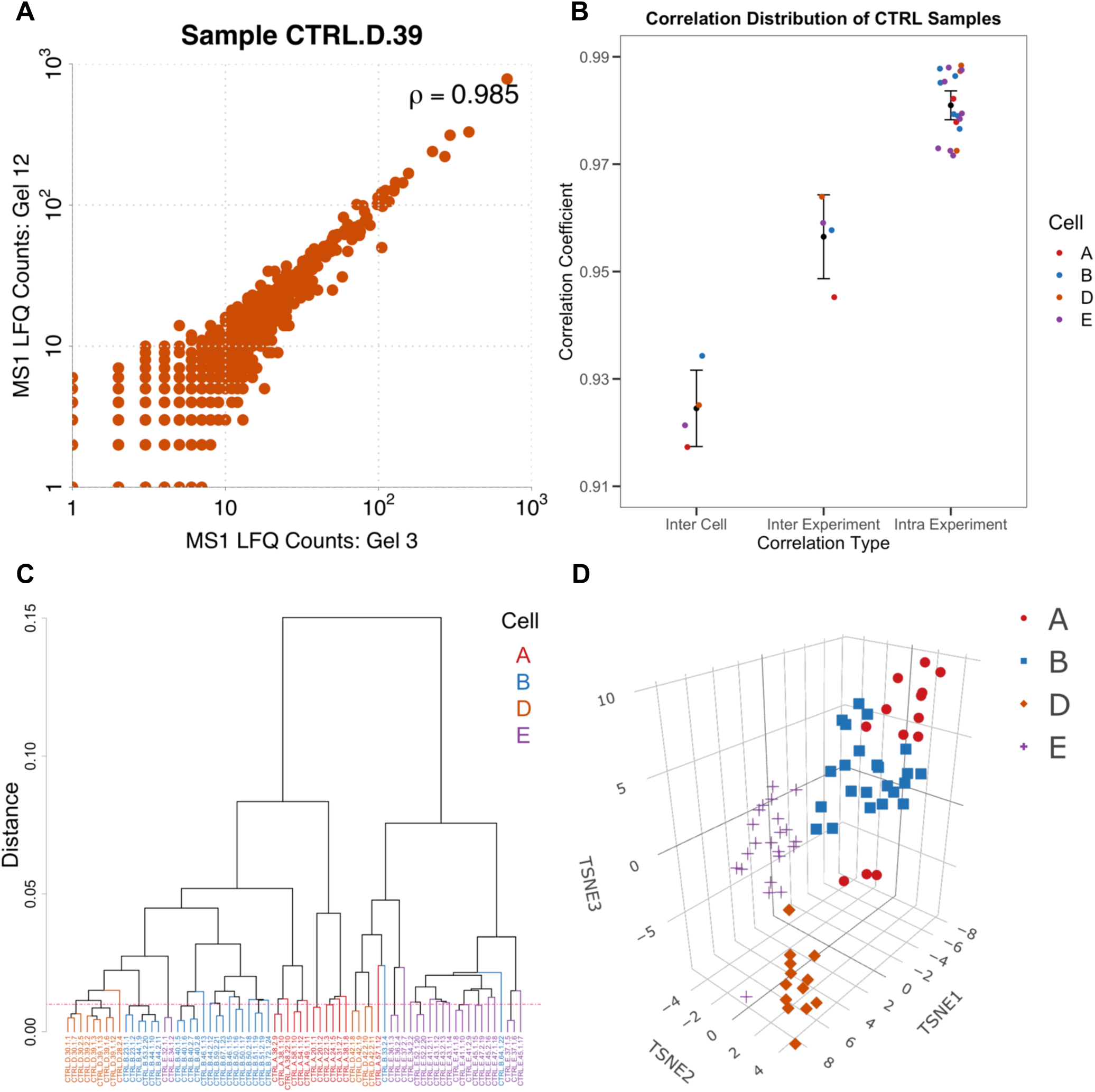
Reproducibility of detected proteins in control samples. **(A)** Normalized MS1 LFQ counts for the same sample measured from two separate SDS-PAGE gels in two different MS/MS runs had strong agreement. **(B)** The correlation (mean and variation) of spectral counts between the control samples from the same experiment for each experiment versus the samples from different experiments in each cell line (cell lines A, B, D, E shown in color coordination). **(C)** The clustering of all biological replicates from four cell lines under control conditions based on the Eucledian distances of spectral counts of replicate samples. **(D)** Accordingly, proteomic signatures of the four cell lines under control conditions across 71 biological replicates showed strong clustering based on the source cell line.

**Figure 4.**
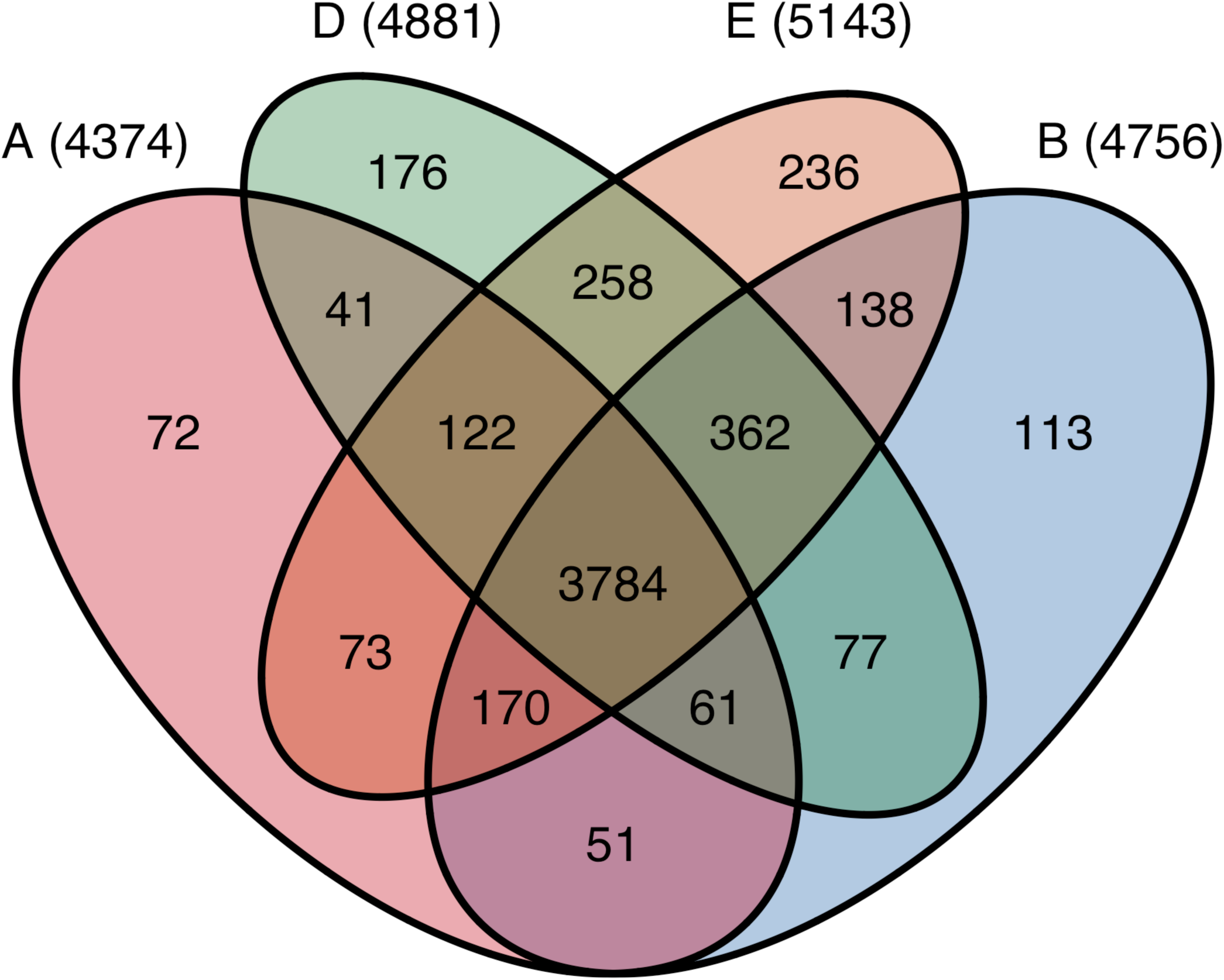
Coverage of identified proteins across cell lines. Venn diagram of the overlapping and unique proteins across all samples for the four PromoCell cardiomyocyte lines (A, B, D, E).

### Identification of differentially expressed proteins

The normalized spectral counts of identified proteins were processed by the edgeR-based computational pipeline to generate differentially expressed proteins (DEPs) associated with statistical significance for each drug treatment in each cell line, as described before ^17^.

### Downstream analysis of differentially expressed proteins

To analyze cell line and drug specific responses, we merged all lists of DEPs of the same cell line treated with the same drug to one new list of DEPs. For each DEP in this new list, we calculated a combined p-value defined as the geometric mean of the individual p-values of the different treatments and a combined log_2_(fold changes) defined as the average mean of the individual log_2_(fold changes). For all downstream analyses, we considered the top 100 DEPs (based on the combined p-values). Combined drug treatments of the different cell lines were hierarchically clustered based on the combined log_2_(fold changes) using Cluster 3.0 ^19,20^. Finally, to investigate the consistency between the different experiments for the same drug treatment across the four cell lines, we ran successive Pearson correlations between different replicates for a given drug treatment of the same cell compared to the correlation among different cell types (**Fig 5A**).

**Figure 5.**
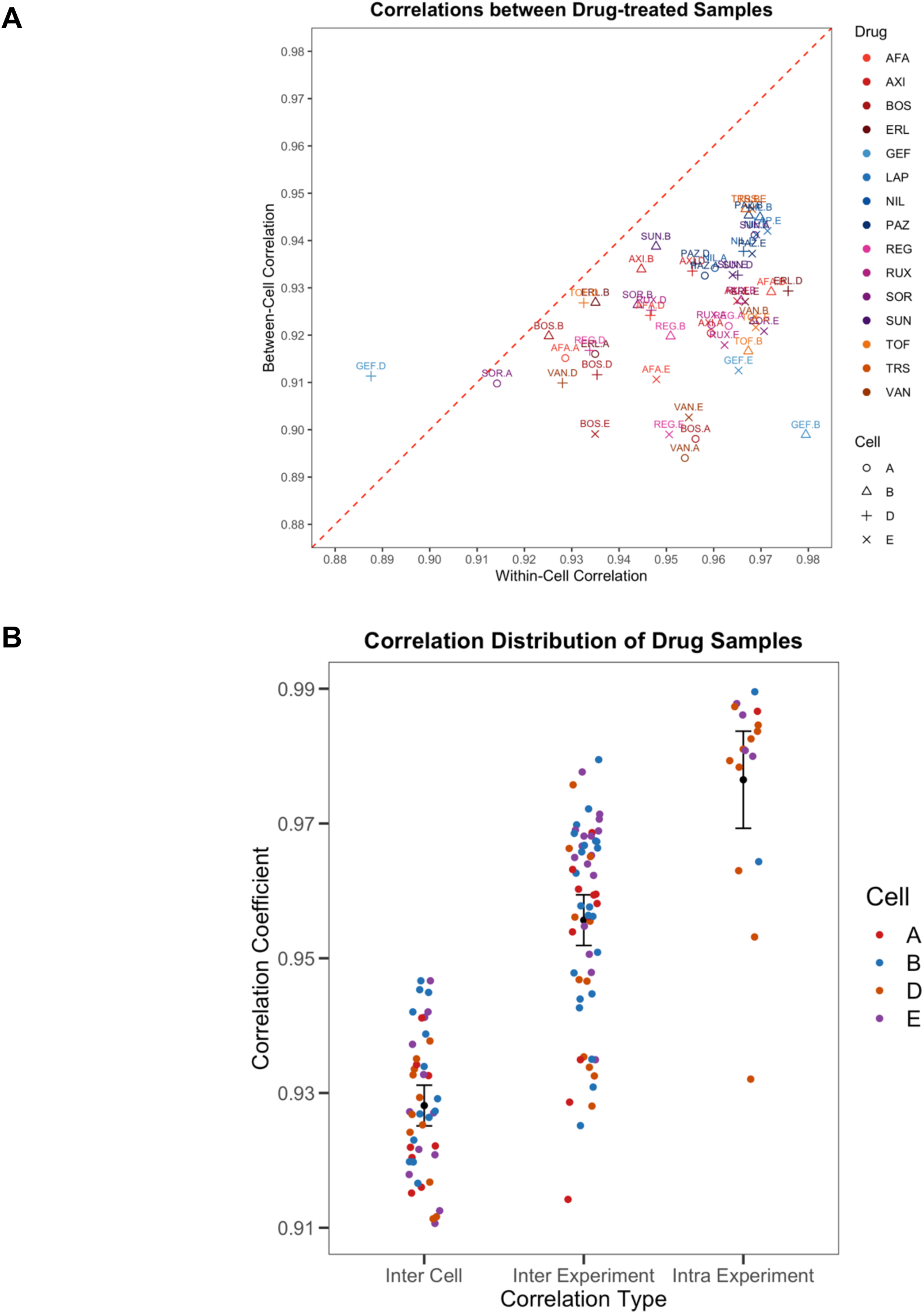
Reproducibility of protein expression measurements for drug-treated samples. **(A)** The similarity and variability of drug-treated samples within and between cell lines as determined by Pearson correlation between drug treated replicates and controls. **(B)** A summary of the drugs showing similar profiles between cells versus those showing different profiles within cells.

## Usage Notes

### Use case 1: Identification of KI drug-induced protein expression profiles

This DToxS dataset can be mined to identify protein expression changes triggered by each KI drug treatment of the four cardiac cell lines derived from human tissues. The proteomic changes identified in this study can be used to understand the cell signal transduction events that may underlie KI-induced cardiotoxicities. Also, with the LC-MS/MS information provided in this dataset, follow-up proteomics assays can be developed to validate the protein expression changes in larger number of samples; such assays may include either targeted assays, *e.g.*, selected or parallel reaction monitoring (SRM or PRM) or untargeted proteomics assays, *e.g.*, data-independent analysis (DIA).

### Use case 2: Comparison of differential KI drug-induced protein expression profiles

This dataset can be analyzed to compare and cluster the proteomic changes induced by different KIs, thus ascertaining the similarities and idiosyncrasies of different drugs and patient (or cell line) specific responses (**Supplementary Fig 3**). One can also quantify the variability and consistency of drug-induced proteomic changes within a given subject (i.e., within a given cell line) compared across a population (i.e., between lines) to determine the generalizability of a given drug effect on cardiomyocytes (**Fig 5B**).

### Use case 3: Integration of transcript and protein expression profiles for comprehensive network studies

Drugs may induce complex changes on either gene or protein expression. Since proteomics and RNAseq changes in complex human cells are not always in agreement, perhaps due to either technical bias, differential regulation or turnover rates; therefore, it may be desirable to integrate both gene and protein expression datasets, in order to obtain more comprehensive understanding of the drug-induced changes among the signaling networks. For example, the DToxS proteomics dataset presented here can be integrated with the DToxS transcriptomics dataset (www.dtoxs.org) to fully understand the effects of KI-induced cardiotoxicities.

### Use case 4: Identification of drug-induced post-translational modifications (PTMs) and regulated proteolysis

Drugs may induce changes among signaling pathways not only *via* the changes of gene and protein expressions, but also *via* the rapid regulations of protein functions by altering specific PTMs (*e.g*., phosphorylation) and regulated proteolytic events (*e.g.* caspase 3 activation) ^24-26^. Therefore, alternative protein database search schemes can be carried out to analyze the raw DToxS proteomics dataset, and identify KI-induced changes among specific PTMs or semi-tryptic or non-tryptic peptides derived from regulated proteolysis (**Supplementary Fig. 4**). Since the current workflow did not include the enrichments of the subproteomes specific for PTMs, the yield for peptides containing PTMs or non-tryptic cleavages will likely be modest. As such, the results from the alternative analysis of the DToxS dataset may provide information for future proteomics studies that focus on the enrichments of specific PTMs, *e.g.* acetylation and phosphorylation.

## Data Citations

1. Xiong et. al (Web link) (2019) (Meta data)
2. Xiong et. al (Web link) (2019) (Raw data)
3. Xiong et. al (Web link) (2019) (Fasta)
4. Xiong et. al (Web link) (2019) (MaxQuant Tables)

## Acknowledgements

The authors are grateful for the support from the NIH Common Fund’s Library of Integrated Cellular Signatures (LINCS) grant U54 HG008098 to R.I., M.R.B., E.A.S. In addition, R.I. is supported in part by NIH GM072853; H.L. is supported in part by NIH R01 GM112415; and E.U.A. is supported in part by NIH R01 DK118222. Mass spectrometer purchases were supported by NIH grant P30 NS046593 from the National Institute of Neurological Disorders and Stroke and another NIH grant 1S10 OD025047 from the Office of the Director of the National Institutes of Health. S.S. and D.V. are supported by NIH U54 HL127624. The content is sole responsibility of the authors and does not necessarily represent the official views of the National Institutes of Health.

## Author Contributions

R.I., E.U.A. and H.L. conceived the project. Y.X. developed the analytical pipeline for proteomic and transcriptomic data processing and tabulated all data and metadata. T.L. executed the LC-MS/MS workflow and data analysis. T.C. performed protein SDS-PAGE analysis and in-gel tryptic digestions. B.H., Y.C., G.J. performed cell culture experiments along with protein and RNA isolation. J.H. developed the bioinformatic pipeline for integrative analysis. S.S., D.V. and J.G. assisted in the development and organization of quality control and metadata standards. H.L. designed the proteomics sample preparation and proteomics analysis workflow from expert input from M.R.B., E.A.S., R.I., and E.U.A. The manuscript was written by H.L., T.L. and E.U.A. with critical input from all the authors. E.U.A. provided overall supervision for all phases of the project.

## Additional Information

Supplemental Information accompanies this paper at http://www.nature.com/sdata

## Supplementary Figures and Tables

**Supplementary Figure 1.**
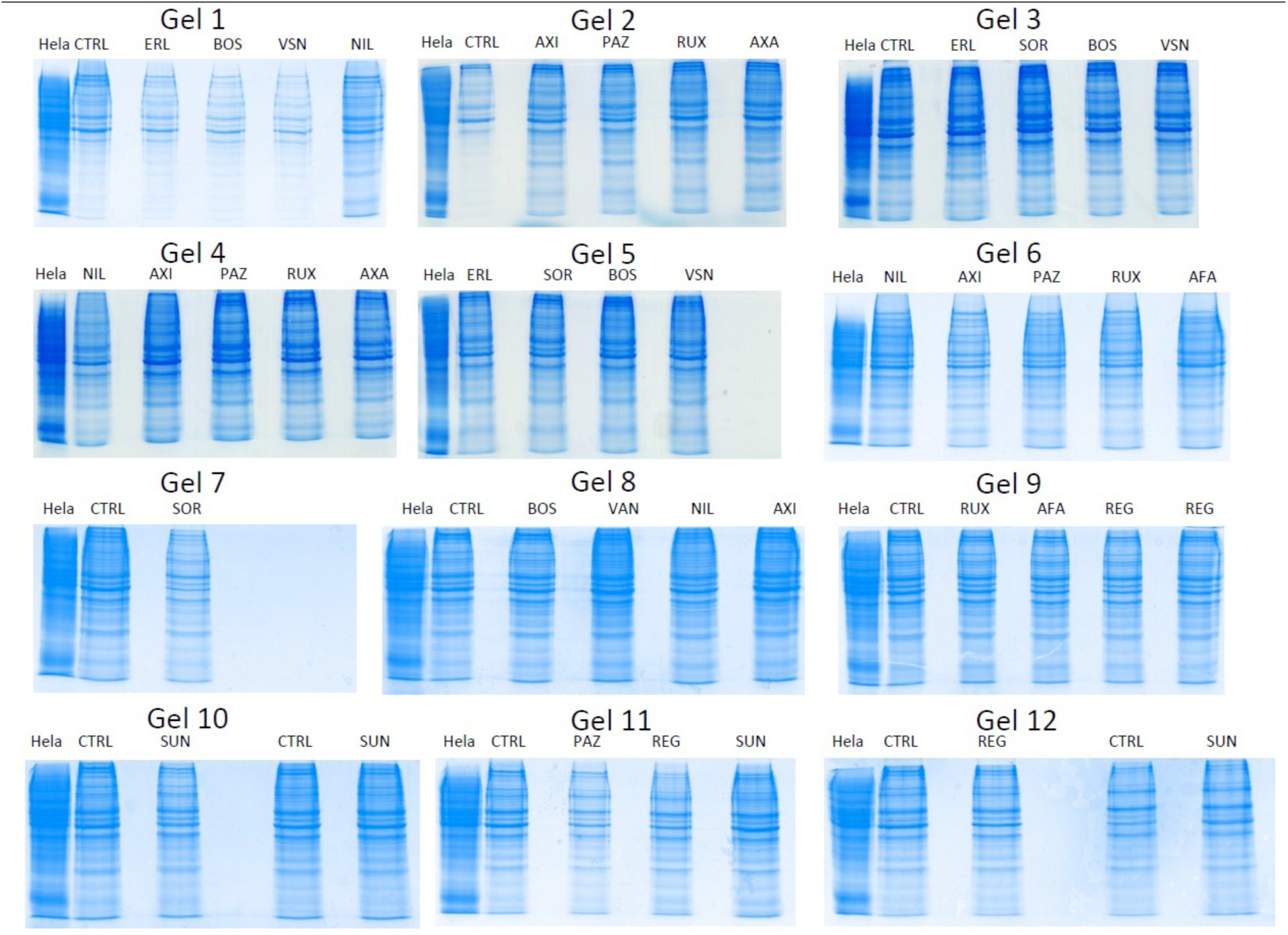
Examples of protein quantification *via* SDS-PAGE. Twenty five micrograms of HeLa cell proteins was loaded along with the proteins recovered from the KI-drug treated cell line A and separated on 10% SDS-PAGE gels. The gel was stained with CBB and the densities of the CBB stain from all lanes are measured. The protein CBB densities in drug-treated samples were compared to that of the HeLa cells, and the protein amounts in each sample was then estimated as described in **Supplementary Table 2**.

**Supplementary Figure 2.**
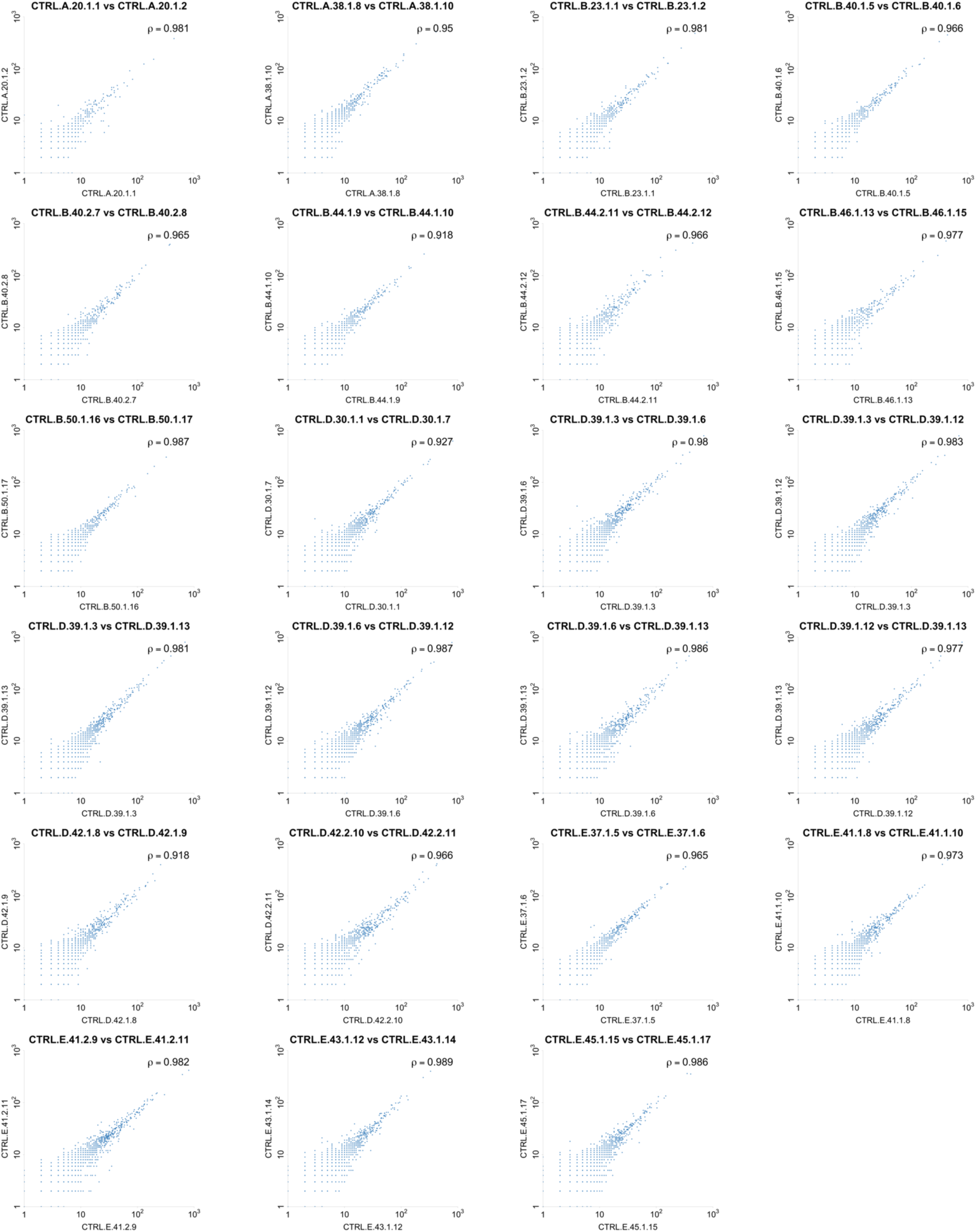
Reproducibility of proteomic quantification for technical replicates. Comparison of normalized MS1 LFQ counts for two technical replicates of the same biological sample that was run on a different SDS-PAGE gel and processed for MS/MS analysis on a different day. Pearson correlation coefficients show strong agreement between multiple runs.

**Supplementary Figure 3.**
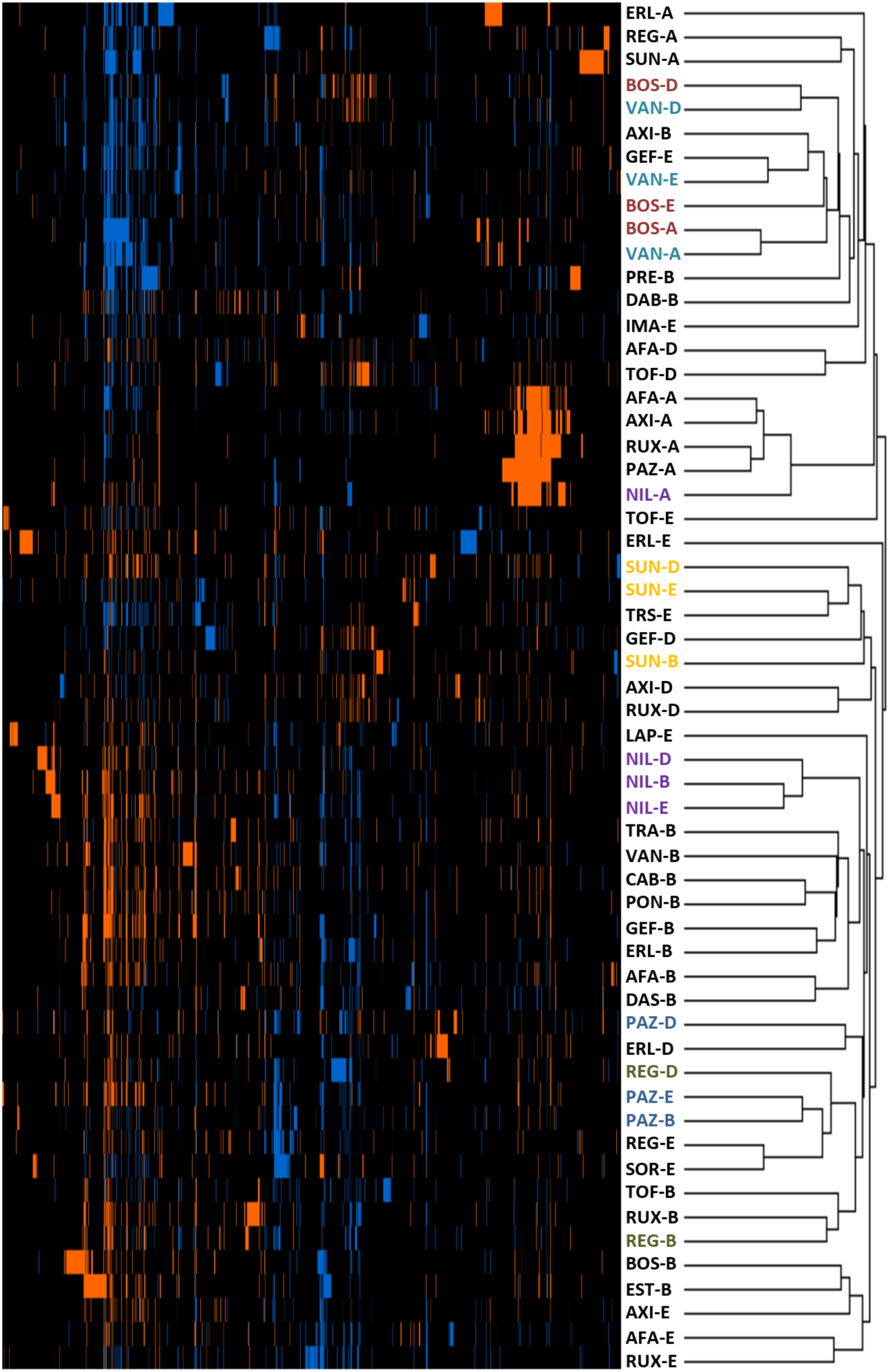
Clustering of KI treated cardiomyocyte cell lines based on top 100 differentially expressed proteins. Cardiomyocyte cell lines treated with KIs that had small proteomic signatures clustered along the cell lines (A, B, D, E); however, those KIs that have large consistent differential proteomic signatures (e.g., SUN, NIL, PAZ, REG) clustered along the drugs regardless of the cell line.

**Supplementary Figure 4.**
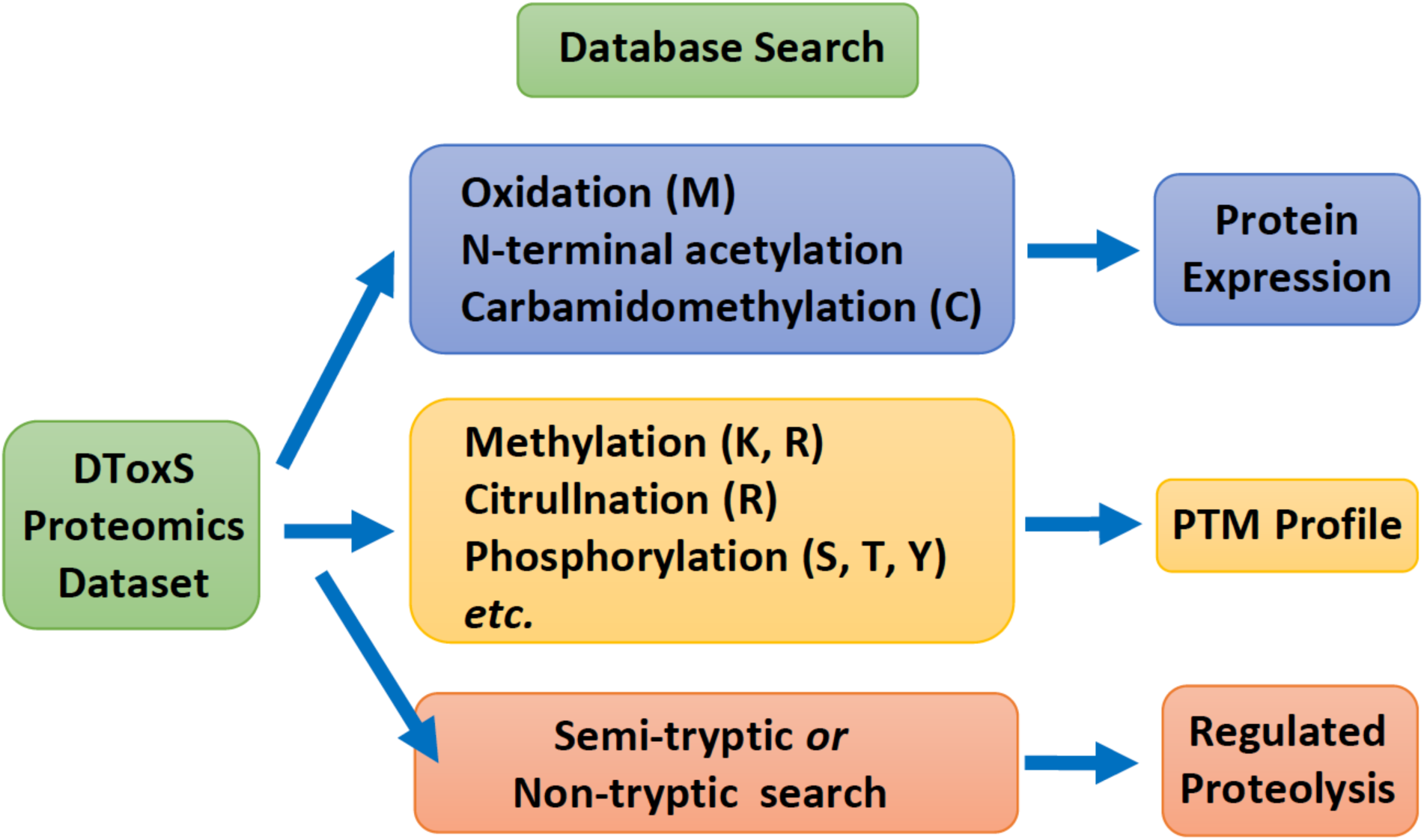
Using the complementary proteomics search methods to identify drug-regulated changes of proteins, PTMs and regulated proteolysis products. The DToxS proteomics dataset can be analyzed with different protein database search methods to obtain different pertinent information, including the changes of protein expression, PTMs and regulated proteolysis of specific peptides.

**Supplementary Table 1.** Drug-treated cell replicates in the DToxS proteomics dataset.

| Drug Treatment | PMC-A | PMC-B | PMC-D | PMC-E |
| --- | --- | --- | --- | --- |
| AFA | 3 | 4 | 4 | 5 |
| AXI | 3 | 4 | 4 | 5 |
| BOS | 5 | 4 | 4 | 5 |
| CAB | 0 | 4 | 0 | 0 |
| DAB | 0 | 4 | 0 | 0 |
| DAS | 0 | 4 | 0 | 0 |
| ERL | 3 | 4 | 4 | 2 |
| EST | 0 | 4 | 0 | 0 |
| GEF | 0 | 4 | 4 | 5 |
| IMA | 0 | 0 | 0 | 3 |
| LAP | 0 | 1 | 0 | 3 |
| PAZ | 3 | 4 | 4 | 5 |
| PON | 0 | 4 | 0 | 0 |
| REG | 4 | 4 | 4 | 5 |
| RUX | 3 | 4 | 4 | 5 |
| SOR | 3 | 3 | 0 | 4 |
| SUN | 3 | 4 | 4 | 6 |
| TOF | 0 | 4 | 4 | 4 |
| TRA | 0 | 4 | 0 | 0 |
| TRS | 0 | 4 | 0 | 4 |
| VAN | 5 | 4 | 4 | 5 |
| VEM | 0 | 3 | 0 | 0 |

\* Four PromoCell cardiomyocyte-like cell lines; 35 separate experiments; 24 drugs
\*\* Control (CTRL) samples: 13 biological replicates for PMC-A, 23 biological replicates for PMC-B, 13 biological replicates for PMC-D, and 22 biological replicates for PMC-E.

**Supplementary Table 2.** Examples of protein quantification *via* SDS-PAGE.

| Gel 1 | Density | Ratio | Protein Amount (μg) | Resuspend Volume(ul) | Conc.(ug/ul) |
| --- | --- | --- | --- | --- | --- |
| HeLa | 4851 | 1 | 25 | 50 | 0.5 |
| CTRL | 3121 | 0.64 | 16 | 32.2 | 0.5 |
| ERL | 2289 | 0.47 | 12 | 23.6 | 0.5 |
| BOS | 2099 | 0.43 | 11 | 21.6 | 0.5 |
| VSN | 2003 | 0.41 | 10 | 20.6 | 0.5 |
| NIL | 3498 | 0.72 | 18 | 36.1 | 0.5 |

| Gel 2 | Density | Ratio | Protein Amount (μg) | Resuspend Volume(ul) | Conc.(ug/ul) |
| --- | --- | --- | --- | --- | --- |
| HeLa | 8468 | 1 | 25 | 50 | 0.5 |
| CTRL | 3622 | 0.43 | 11 | 21.4 | 0.5 |
| AXI | 5688 | 0.67 | 17 | 33.6 | 0.5 |
| PAZ | 5929 | 0.7 | 18 | 35 | 0.5 |
| RUX | 6428 | 0.76 | 19 | 38 | 0.5 |
| AFA | 7152 | 0.84 | 21 | 42.2 | 0.5 |

| Gel 3 | Density | Ratio | Protein Amount (μg) | Resuspend Volume(ul) | Conc.(ug/ul) |
| --- | --- | --- | --- | --- | --- |
| HeLa | 8468 | 1 | 25 | 50 | 0.5 |
| CTRL | 3622 | 0.43 | 11 | 21.4 | 0.5 |
| AXI | 5688 | 0.67 | 17 | 33.6 | 0.5 |
| PAZ | 5929 | 0.7 | 18 | 35 | 0.5 |
| RUX | 6428 | 0.76 | 19 | 38 | 0.5 |
| AFA | 7152 | 0.84 | 21 | 42.2 | 0.5 |

| Gel 4 | Density | Ratio | Protein Amount (μg) | Resuspend Volume(ul) | Conc.(ug/ul) |
| --- | --- | --- | --- | --- | --- |
| HeLa | 9789 | 1 | 25 | 50 | 0.5 |
| NIL | 5914 | 0.6 | 15 | 30.2 | 0.5 |
| AXI | 7570 | 0.77 | 19 | 38.7 | 0.5 |
| PAZ | 7507 | 0.77 | 19 | 38.3 | 0.5 |
| RUX | 7172 | 0.73 | 18 | 36.6 | 0.5 |
| AFA | 6802 | 0.69 | 17 | 34.7 | 0.5 |

| Gel 5 | Density | Ratio | Protein Amount (μg) | Resuspend Volume(ul) | Conc.(ug/ul) |
| --- | --- | --- | --- | --- | --- |
| HeLa | 8720 | 1 | 25 | 50 | 0.5 |
| ERL | 5983 | 0.69 | 17 | 34.3 | 0.5 |
| SOR | 6080 | 0.7 | 18 | 34.9 | 0.5 |
| BOS | 6559 | 0.75 | 19 | 37.6 | 0.5 |
| VSN | 6436 | 0.74 | 19 | 36.9 | 0.5 |

| Gel 6 | Density | Ratio | Protein Amount (μg) | Resuspend Volume(ul) | Conc.(ug/ul) |
| --- | --- | --- | --- | --- | --- |
| HeLa | 2337 | 1 | 25 | 50 | 0.5 |
| NIL | 1988 | 0.85 | 21 | 42.5 | 0.5 |
| AXI | 1660 | 0.71 | 18 | 35.5 | 0.5 |
| PAZ | 1844 | 0.79 | 20 | 39.5 | 0.5 |
| RUX | 1757 | 0.75 | 19 | 37.6 | 0.5 |
| AFA | 1895 | 0.81 | 20 | 40.5 | 0.5 |

| Gel 7 | Density | Ratio | Protein Amount (μg) | Resuspend Volume(ul) | Conc.(ug/ul) |
| --- | --- | --- | --- | --- | --- |
| HeLa | 3409 | 1 | 25 | 50 | 0.5 |
| CTRL | 2767 | 0.81 | 20 | 40.6 | 0.5 |
| SOR | 2006 | 0.59 | 15 | 29.4 | 0.5 |

| Gel 8 | Density | Ratio | Protein Amount (μg) | Resuspend Volume(ul) | Conc.(ug/ul) |
| --- | --- | --- | --- | --- | --- |
| HeLa | 2439 | 1 | 25 | 50 | 0.5 |
| CTRL | 1786 | 0.73 | 18 | 36.6 | 0.5 |
| BOS | 1878 | 0.77 | 19 | 38.5 | 0.5 |
| VAN | 2200 | 0.9 | 23 | 45.1 | 0.5 |
| NIL | 1953 | 0.8 | 20 | 40 | 0.5 |
| AXI | 2096 | 0.86 | 22 | 43 | 0.5 |

| Gel 9 | Density | Ratio | Protein Amount (μg) | Resuspend Volume(ul) | Conc.(ug/ul) |
| --- | --- | --- | --- | --- | --- |
| HeLa | 2708 | 1 | 25 | 50 | 0.5 |
| CTRL | 2282 | 0.84 | 21 | 42.1 | 0.5 |
| RUX | 2144 | 0.79 | 20 | 39.6 | 0.5 |
| AFA | 2199 | 0.81 | 20 | 40.6 | 0.5 |
| REG | 2965 | 1.09 | 27 | 54.7 | 0.5 |
| REG | 2070 | 0.76 | 19 | 38.2 | 0.5 |

| Gel 10 | Density | Ratio | Protein Amount (μg) | Resuspend Volume(ul) | Conc.(ug/ul) |
| --- | --- | --- | --- | --- | --- |
| HeLa | 2869 | 1 | 25 | 50 | 0.5 |
| CTRL | 2236 | 0.78 | 20 | 39 | 0.5 |
| SUN | 1880 | 0.66 | 17 | 32.8 | 0.5 |
| CTRL | 2262 | 0.79 | 20 | 39.4 | 0.5 |
| SUN | 2240 | 0.78 | 20 | 39 | 0.5 |

| Gel 11 | Density | Ratio | Protein Amount (μg) | Resuspend Volume(ul) | Conc.(ug/ul) |
| --- | --- | --- | --- | --- | --- |
| HeLa | 2317 | 1 | 25 | 50 | 0.5 |
| CTRL | 1760 | 0.76 | 19 | 38 | 0.5 |
| PAZ | 1321 | 0.57 | 14 | 28.5 | 0.5 |
| REG | 1575 | 0.68 | 17 | 34 | 0.5 |
| SUN | 1888 | 0.81 | 20 | 40.7 | 0.5 |

| Gel 12 | Density | Ratio | Protein Amount (μg) | Resuspend Volume(ul) | Conc.(ug/ul) |
| --- | --- | --- | --- | --- | --- |
| HeLa | 1444 | 1 | 25 | 50 | 0.5 |
| CTRL | 1121 | 0.78 | 20 | 38.8 | 0.5 |
| REG | 995 | 0.69 | 17 | 34.5 | 0.5 |
| CTRL | 1062 | 0.74 | 19 | 36.8 | 0.5 |
| SUN | 1023 | 0.71 | 18 | 35.4 | 0.5 |

The CBB protein density of the entire lane for each sample lane was measured using ImageJ software (NIH), and the protein amount for each drug-treated sample was calculated using the following equation: Protein amount = HeLa cell protein amount × (Sample density / HeLa cell density). After in-gel tryptic digestions, the peptides were dissolved in HPLC solvents to obtain a final concentration of 0.5 μg/mL, based on the calculated protein amounts.

